# Lipid interactions enhance activation and potentiation of cystic fibrosis transmembrane conductance regulator (CFTR)

**DOI:** 10.1101/495010

**Authors:** Stephanie Chin, Mohabir Ramjeesingh, Maurita Hung, June Ereño-Oreba, Chris Ing, Zhi W. Zeng, Hong Cui, Régis Pomès, Jean-Philippe Julien, Christine E. Bear

**Affiliations:** Programme in Molecular Medicine, Research Institute, Hospital for Sick Children, Toronto, ON, Canada; Department of Biochemistry, University of Toronto, Toronto, ON, Canada; Department of Physiology, University of Toronto, Toronto, ON, Canada; Department of Immunology, University of Toronto, Toronto, ON, Canada

**Author notes:** Corresponding Author: Christine E. Bear Programme in Molecular Medicine, Peter Gilgan Centre for Research and Learning, Hospital for Sick Children, 686 Bay Street, Room 20.9706, Toronto, ON, M5G 0A4, Canada.

## Abstract

The recent cryo-electron microscopy structures of phosphorylated, ATP-bound CFTR in detergent micelles failed to reveal an open anion conduction pathway as expected on the basis of previous functional studies in biological membranes. We tested the hypothesis that interaction of CFTR with lipids is important for opening of its channel. Interestingly, molecular dynamics studies revealed that phospholipids associate with regions of CFTR proposed to contribute to its channel activity. More directly, we found that CFTR purified together with associated lipids using the amphipol: A8-35, exhibited higher rates of catalytic activity, channel activation and potentiation using ivacaftor, than did CFTR purified in detergent. Catalytic activity in CFTR detergent micelles was partially rescued by addition of phospholipids plus cholesterol, arguing that these lipids contribute directly to its modulation. In summary, these studies highlight the importance of lipids in regulated CFTR channel activation and potentiation.

## Introduction

Cystic Fibrosis is caused by mutations of the Cystic Fibrosis Transmembrane conductance Regulator (*CFTR*) gene that codes for the ATP-binding cassette (ABC) anion channel, CFTR (Tsui, Buchwald et al. 1985, Kerem, Rommens et al. 1989, Kartner, Hanrahan et al. 1991). CFTR channel activity and anion conduction through its membrane pore is regulated by protein kinase A (PKA) phosphorylation of its regulatory (R) domain as well as ATP binding and hydrolysis by its nucleotide binding domains (NBDs) (Chang, Tabcharani et al. 1993, Winter and Welsh 1997, Seibert, Chang et al. 1999, Ramjeesingh, Ugwu et al. 2008, Hwang and Sheppard 2009, Mihalyi, Torocsik et al. 2016). Biochemical and electrophysiological studies support a model where phosphorylation promotes dissociation of the R domain from inhibitory locations at the interface between the NBDs to facilitate their dimerization and ATPase activity (Kogan, Ramjeesingh et al. 2001, Kogan, Ramjeesingh et al. 2002, Kidd, Ramjeesingh et al. 2004, Vergani, Lockless et al. 2005, Aleksandrov, Aleksandrov et al. 2007, Baker, Hudson et al. 2007, Stratford, Ramjeesingh et al. 2007, Ramjeesingh, Ugwu et al. 2008, Hwang and Sheppard 2009, Kanelis, Hudson et al. 2010, Dawson, Farber et al. 2013). This model predicts that the NBDs of the unphosphorylated and nonconductive form of CFTR may be dissociated, whereas the NBDs will be dimerized in the phosphorylated and conductive form of the protein.

Recently, four structures of CFTR have been determined using cryo-electron microscopy (EM) at resolutions of 3.2 – 3.9 Å (Zhang and Chen 2016, Liu, Zhang et al. 2017, Zhang, Liu et al. 2017, Zhang, Liu et al. 2018). These structures include the unphosphorylated, ATP-free and phosphorylated, ATP-bound forms and the unphosphorylated, ATP-free form of the zCFTR and hCFTR proteins (Zhang and Chen 2016, Liu, Zhang et al. 2017, Zhang, Liu et al. 2017, Zhang, Liu et al. 2018). Comparison of the structures show major conformational changes upon PKA phosphorylation and ATP binding that are consistent with the molecular mechanism proposed for opening of the conduction pore through CFTR. Such changes included the loss of stable R domain interactions at the cleft between the two transmembrane domain (TMD)-NBD halves of the CFTR molecule and tighter association of the NBDs (Zhang and Chen 2016, Zhang, Liu et al. 2017, Zhang, Liu et al. 2018). There were also subtle conformational changes at the transmembrane segments (TMs), notably at TM8 and TM12, upon PKA phosphorylation and ATP binding (Zhang, Liu et al. 2017, Zhang, Liu et al. 2018). These structures suggest that both TM8 and TM12 helices rotate to participate in novel interactions upon CFTR phosphorylation (Zhang, Liu et al. 2017, Zhang, Liu et al. 2018). Unexpectedly, the structure of phosphorylated, ATP-bound CFTR proteins did not reveal a continuous pathway for anion conduction (Zhang, Liu et al. 2017, Zhang, Liu et al. 2018). Hence, factors required to stabilize the open conduction pathway may be lacking in the CFTR proteins studied by cryo-EM.

Recent studies have shown that ABC transporters specifically bind to lipids and these protein: lipid interactions are structurally important (Taylor, Manolaridis et al. 2017, Jackson, Manolaridis et al. 2018, Josts, Nitsche et al. 2018). Phospholipids and cholesterol are bound to ABCG2 in a recent cryo-EM structure (Taylor, Manolaridis et al. 2017, Jackson, Manolaridis et al. 2018) and conformers of MsbA solved in lipid containing nanodiscs differ from those determined in detergent micelles (Josts, Nitsche et al. 2018). Furthermore, lipid interactions are known to modify the activity of several ABC transporters including: P-glycoprotein (Ambudkar, Lelong et al. 1998, Ritchie, Grinkova et al. 2009), ABCG1 (Hirayama, Kimura et al. 2013), LmrA (Infed, Hanekop et al. 2011) and MsbA (Doerrler and Raetz 2002, Kawai, Caaveiro et al. 2011). In fact, the addition of lipids was found to stabilize CFTR after its purification in detergent and modulate its ATPase activity (Yang, Wang et al. 2014, Hildebrandt, Khazanov et al. 2017). Hence, we hypothesized that the conductance competent form of phosphorylated, ATP-bound CFTR protein is stabilized through lipid interactions. To test this idea, we developed a novel method to purify CFTR in a detergent-free environment. We confirmed that, relative to detergent-purified protein, CFTR extracted from mammalian membranes using amphipol A8-35 retains associated lipids as well as exhibits enhanced ATPase activity, channel function and response to the first drug targeting mutant CFTR.

## Results

### Molecular dynamics (MD) simulation studies of “pre-open” state of CFTR reveal regions with propensity for specific phospholipid interaction

We performed multiple microsecond-long MD simulations of a homology model of hCFTR in the “pre-open” state (phosphorylated, ATP-bound) embedded in 1,2-palmitoyl-oleoyl-sn-glycero-3-phosphocholine (POPC) or 1,2-palmitoyl-oleoyl-sn-glycero-3-phosphoserine (POPS) bilayers, two lipids shown previously to support different levels of catalytic activity by CFTR (Hildebrandt, Khazanov et al. 2017). By calculating the number density of POPC (Figure 1a, green volumes) or POPS (Figure 1b, blue volumes), we observe an annular lipid shell (transparent surface) wrapping around the TM region of hCFTR, revealing multiple sites enriched for protein: lipid interactions (solid shapes). The number of lipid sites within the annulus is 64 ± 3 for POPC (Figure 1a, transparent green surface) and 70 ± 4 for POPS (Figure 1b, transparent blue surface) bilayers, respectively (based on heavy-atom contacts within 4.5 Å). Across multiple simulation repeats, we observe lipid binding sites that are enriched for interactions with a particular lipid species. We show a differential association of POPC over POPS with the outer vestibule of the CFTR protein (Figure 1c, green volumes) and POPS over POPC with the inner vestibule of the channel (Figure 1d, blue volumes). POPC has a propensity to associate with a groove between TM7 and TM9 in the outer vestibule. There are multiple regions for which there is an increased propensity for POPS to associate with CFTR in the inner membrane leaflet, including a cleft between TM4 and TM6 and another groove between TM10 and TM11. As yet, the molecular basis for the positive effect of POPS on CFTR ATPase activity remains unknown (Hildebrandt, Khazanov et al. 2017), but our analyses support the hypothesis that its interaction with positively charged residues lining the cleft between TM4 and TM6 may disrupt previously described interactions with organic anions known to be inhibitory to CFTR ATPase activity (Kogan, Ramjeesingh et al. 2001, El Hiani and Linsdell 2015, El Hiani, Negoda et al. 2016). Overall, these relative propensity maps of lipid interaction with the “pre-open” state of CFTR correlate with regions that have been identified as important for channel opening (Wang, El Hiani et al. 2014, El Hiani and Linsdell 2015, El Hiani, Negoda et al. 2016, Hwang, Yeh et al. 2018).

**Figure 1:**
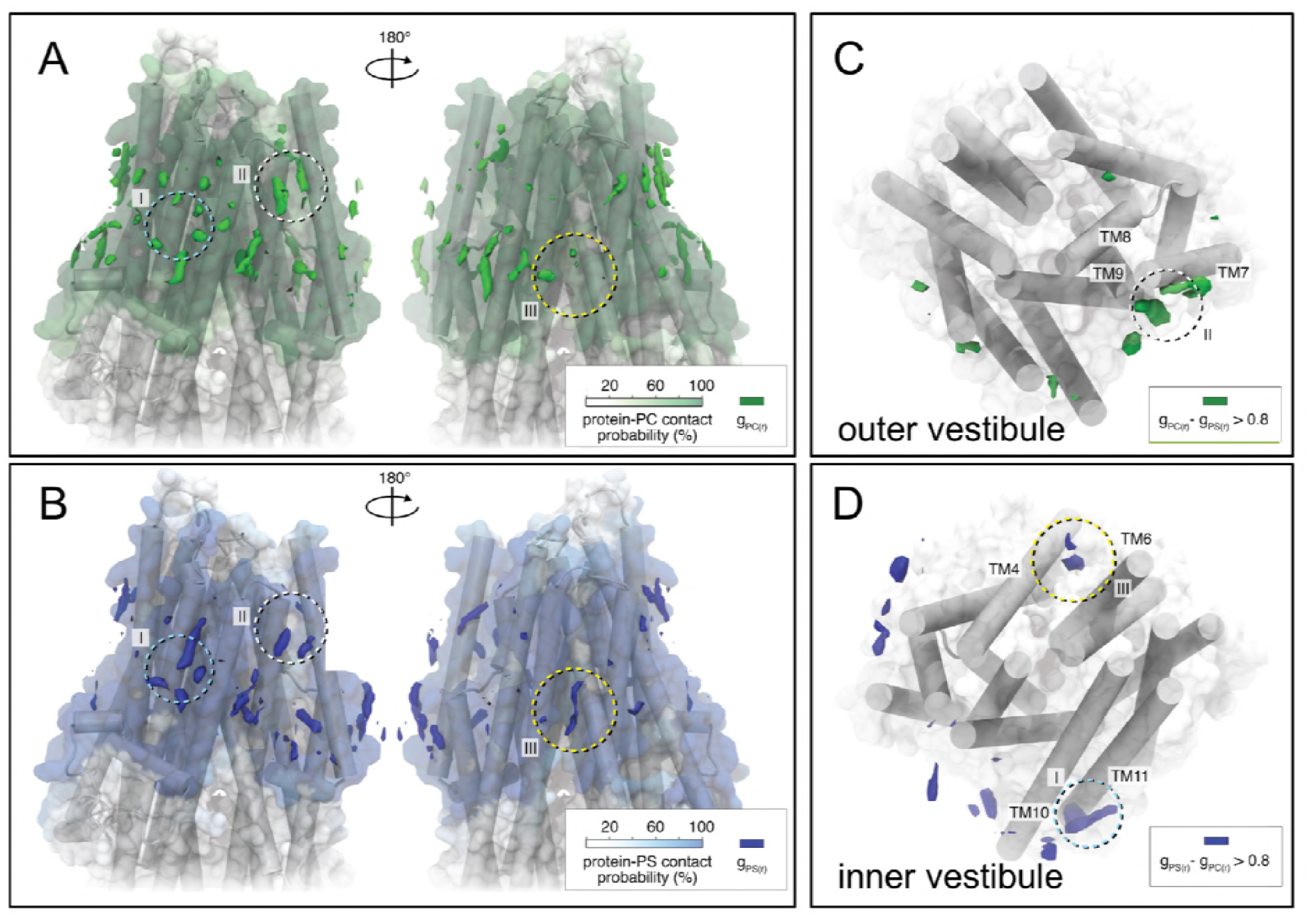
Phospholipids interact with CFTR. (a and b) Molecular renderings of hCFTR models viewed from a side orientation. The spatial distribution function of lipid heavy atoms over all MD simulations is depicted as colored volumes for (a) POPC (g_PC(r)_, green) and (b) POPS (g_PS(r)_, blue). Each map is normalized by the average number density of the lipids in a neat bilayer and only regions with density > 160% of this value are shown. A transparent surface is colored by the percentage of simulation data where heavy-atom contacts are made between protein and lipids atoms. A difference map shows regions with increased binding for (c) POPC (g_PC(r)_ - g_PS(r)_ > 0.8) in the outer leaflet and (d) POPS (g_PS(r)_ - g_PC(r)_ > 0.8) in the inner leaflet. Labels I, II, and III highlight regions where differences can be observed between these spatial distribution functions. Preferential binding of POPC was found at the external cleft between TM7 and TM9 (region II). Preferential binding of POPS was found at the internal clefts between TM10 and TM11 (region I) and between TM4 and TM6 (region III).

### CFTR: lipid complexes are purified using amphipol A8-35

Full-length human Wt-CFTR, bearing its native sequence except for two affinity tags, a FLAG tag (on the amino terminus) and a 10x histidine-tag (on the carboxy terminus) was expressed in HEK293F cells. Amphipol A8-35 at 30 mg was effective in solubilizing CFTR from 1 ml of 2 mg/ml final concentration of resuspended membranes, which corresponds to an amphipol concentration of 3% (w/v) (Figure 2a, lane 1). Finally, CFTR was purified to near homogeneity in a single step by affinity to anti-FLAG^®^ magnetic beads (Figure 2a, lane 3 and 4). In SDS-PAGE, purified CFTR migrated as expected for the mature, complex glycosylated band C of the protein (170-180 kDa, Figure 2a, lane 5). The immature, core glycosylated band B of CFTR was also present in the eluted sample as a band at around 140-150 kDa (Figure 2a, lane 5).

**Figure 2:**
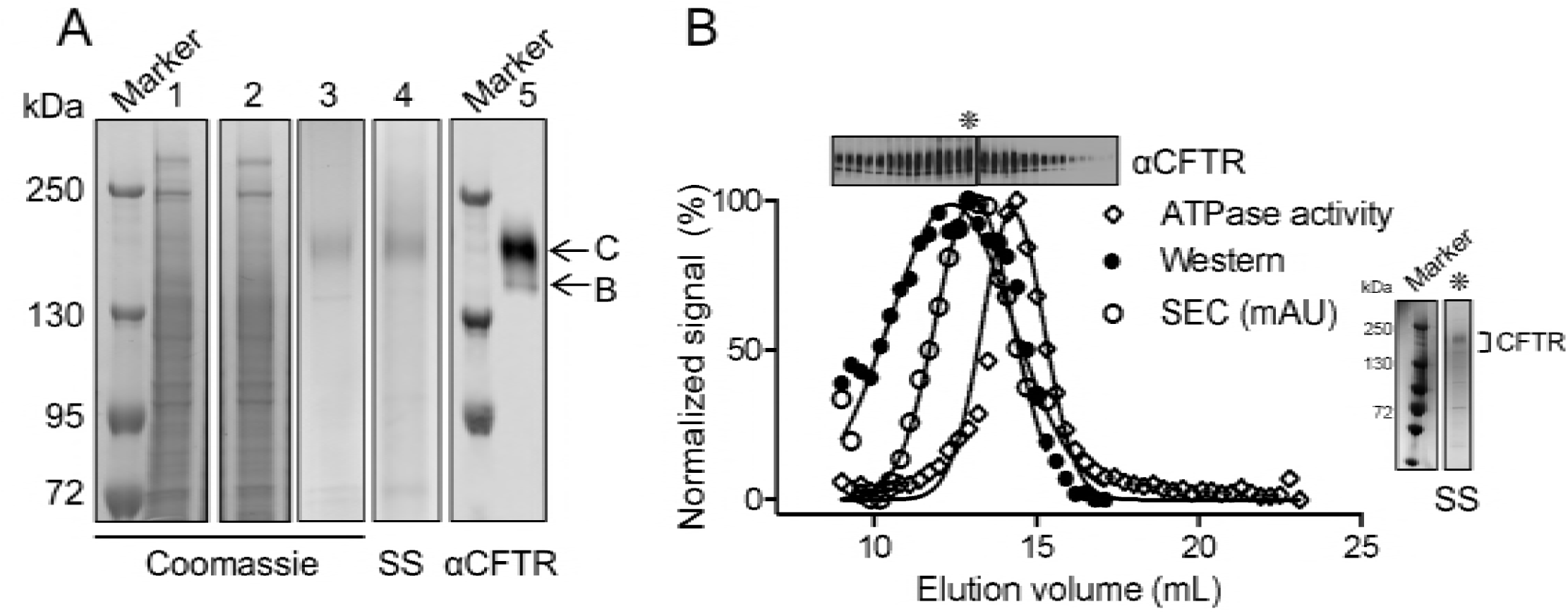
Functional CFTR can be purified using amphipol. (a) Protein gels stained by Coomassie blue of lysate solubilized by amphipol (1), unbound fraction (2) and eluate from anti-FLAG^®^ M2 magnetic beads (3). Eluate from anti-FLAG^®^ M2 magnetic beads was also stained by silver stain (SS) (4) and immunoblotted (5) showing that CFTR was effectively purified as a relatively pure population. CFTR appeared as two bands that include a band at approximately 170-180 kDa that represents the mature, complex glycosylated form of the protein (band C) and a band at approximately 140-150 kDa that represents the immature, core glycosylated form of the protein (band B). (b) Overlay of normalized signal of CFTR ATPase activity (open diamonds) and CFTR protein detected by immunoblot (closed circles) to SEC trace on a Superose 6 10/300 GL (GE Healthcare) (open circles) showing that the SEC peak at (asterisk) corresponds to CFTR protein separation and the shoulder of that peak corresponds to functional CFTR. SS of the SEC peak (asterisk) shows the purified CFTR protein is relatively pure.

Size exclusion chromatography (SEC) was used to characterize the purified CFTR protein on the basis of differences in size using 280 nm to monitor the elution of CFTR (Figure 2b). The fractions eluting between elution volume of 9-17 ml containing both core and complex glycosylated CFTR, as confirmed by silver staining and immunoblotting, co-eluted with ATPase activity (Figure 2b). Interestingly, the peak ATPase activity at elution volume of 14.5 ml was slightly displaced from the peak CFTR protein abundance at elution volume of 13 ml, suggesting that the population of purified CFTR molecules is heterogeneous with respect to specific enzyme activity.

Previous studies found that amphipols do not compete with lipids, which remain associated to membrane protein: amphipol complexes (Martinez, Gohon et al. 2002, Gohon, Dahmane et al. 2008, Bechara, Bolbach et al. 2012). Thus, we were interested in determining whether CFTR purified using amphipol also retained lipids. We compared the amphipol-based purification to that employed by the Chen group in their cryo-EM studies (Zhang and Chen 2016, Liu, Zhang et al. 2017, Zhang, Liu et al. 2017). First, we confirmed that CFTR solubilized in Lauryl Maltose Neopentyl Glycol (LMNG) detergent micelles (Zhang and Chen 2016, Liu, Zhang et al. 2017, Zhang, Liu et al. 2017) could be purified using the same protocol that we developed for the amphipol preparation (Figure 3a,b). Then, the relative degree of phospholipid association was compared in the amphipol- and detergent-based purifications. Choline-containing lipids (i.e. phosphatidylcholine or PC, lecithin, lysolecithin, and sphingomyelin) were quantified using a fluorimetric assay after normalization for CFTR protein abundance (Refer to *CFTR quantification* section in Materials and Methods for more details on the method). We detected a higher number of approximately 56 to 76 choline-containing phospholipids associated per CFTR molecule in amphipol compared to approximately 5 to 12 phospholipids associated per CFTR in LMNG detergent micelles (Figure 3c). Interestingly, the number of measured lipid molecules correlated with the number determined in our MD simulations of POPC or POPS. Notably, the lipids that associated with CFTR in the amphipol purification include cholesterol, PC and phosphatidylethanolamine (PE) as studied by lipid thin-layer chromatography (TLC) (Figure 3d). We cannot exclude the presence of additional lipids, given the resolution of the solvent system employed for TLC. In fact, we expect that phosphatidylserine (PS) is also associated given its dominant presence in the inner leaflet of biological membranes.

**Figure 3:**
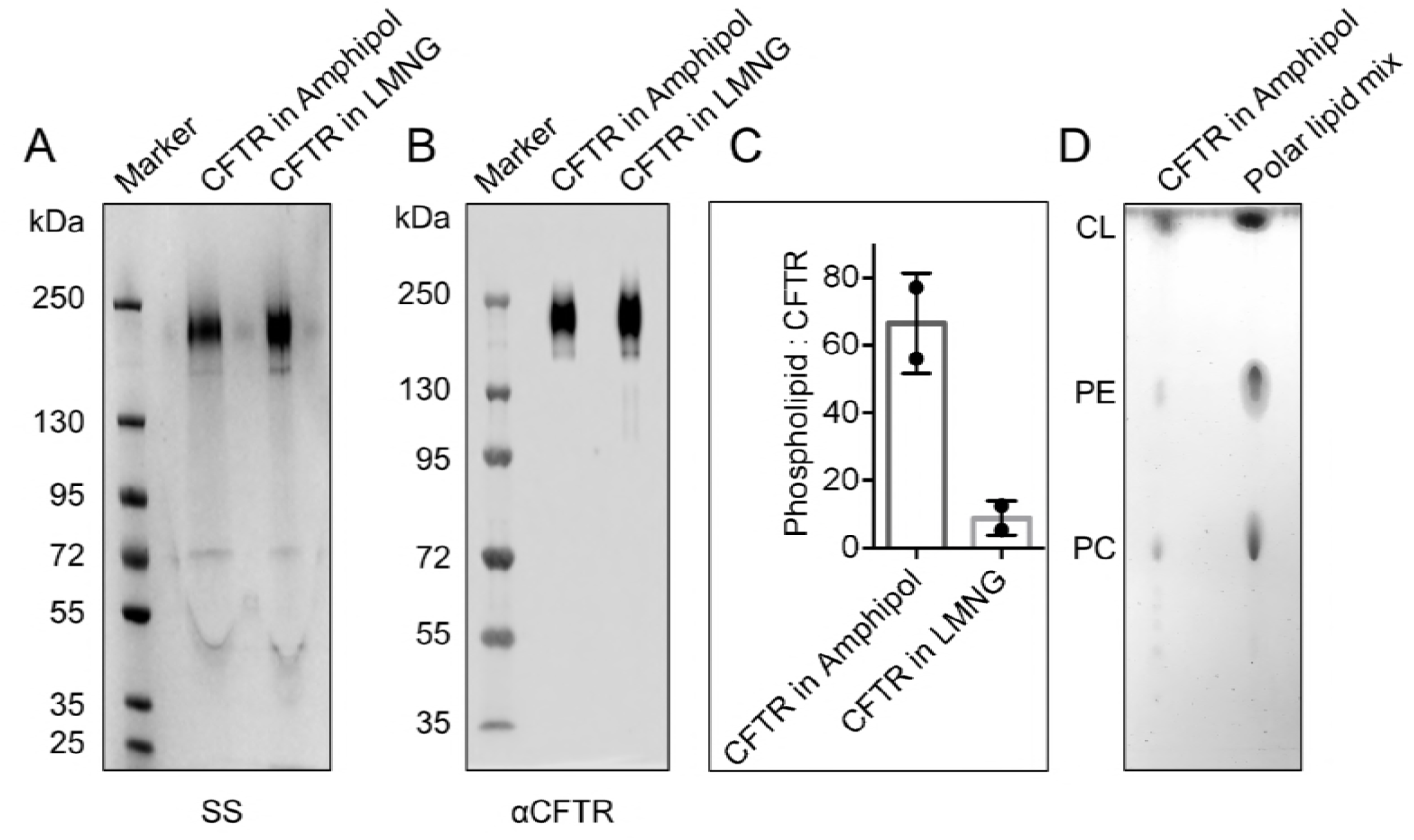
Yield of CFTR is similar for amphipol and LMNG purifications and amphipol purified CFTR retains an annulus of phospholipids and cholesterol. (a) SS and (b) immunoblot of purified CFTR from amphipol and LMNG purifications showing that both purifications yielded high purity of CFTR protein at similar yields. (c) Phospholipid analysis showing that purified CFTR solubilized in amphipol contained more phospholipids (56 to 76 phospholipids per CFTR molecule) than CFTR solubilized in LMNG (5 to 12 phospholipids per CFTR molecule) after normalizing to protein amounts. The data is presented as a range of n = 2 technical replicates. (d) Lipid TLC showing that extracted lipids from purified CFTR in amphipol include cholesterol, PE and PC as confirmed by similar migration of known lipids from polar lipid mix standard. These lipids were not present in amphipol by itself.

### CFTR: lipid: amphipol complexes exhibit higher specific ATPase activity and higher functional reconstitution as regulated anion channels than CFTR: detergent complexes

We compared the specific ATPase activities of CFTR purified using amphipol or LMNG after PKA phosphorylation (P) to ensure that both preparations were maximally phosphorylated, a modification known to be important for CFTR function as an enzyme and a channel (Chang, Tabcharani et al. 1993, Li, Ramjeesingh et al. 1996, Winter and Welsh 1997, Seibert, Chang et al. 1999, Szellas and Nagel 2003). We found that the K_m_ values of ATP were similar for the two samples: K_m_ of 0.27 ± 0.08 mM ATP for the purified P-CFTR in amphipol and K_m_ of 0.32 ±?0.14 mM ATP for the purified P-CFTR in LMNG detergent micelles (Figure 4a). Interestingly, the maximal ATPase activities, after normalization to protein amounts (Refer to *CFTR quantification* section in Materials and Methods for more details on the method), were significantly higher in the purified P-CFTR: amphipol complexes than in the purified P-CFTR: LMNG micelles (V_max_ of 23.90 ± 1.91 nmol phosphate/mg protein/min versus V_max_ of 5.54 ± 0.66 nmol phosphate/mg protein/min respectively, Figure 4a).

**Figure 4:**
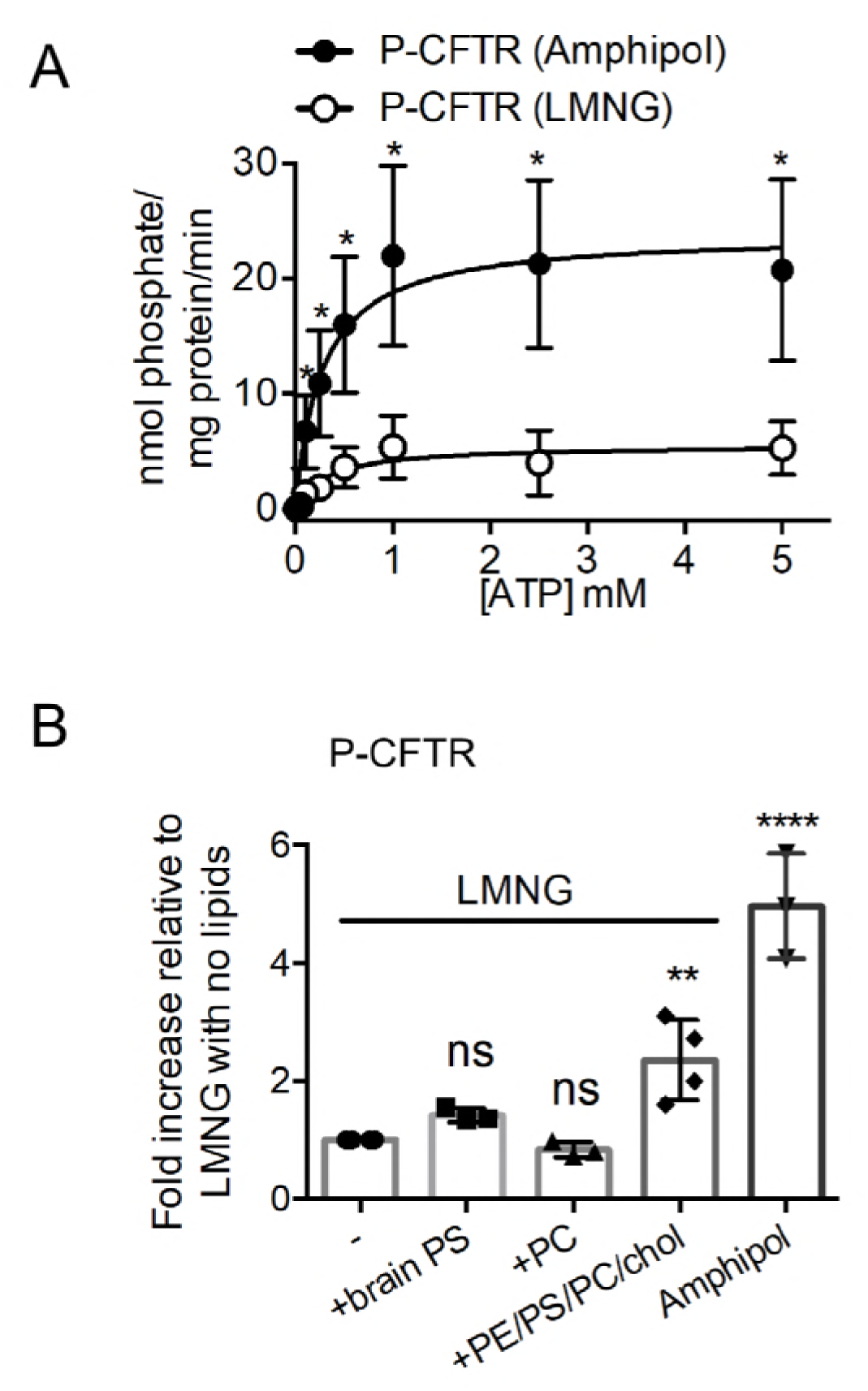
Specific ATPase activity of phosphorylated CFTR is higher in amphipol preparation than in detergent preparation and this difference is related to lipid associations of CFTR. (a) ATPase activity of P-CFTR in amphipol (closed circles) and in LMNG detergent (open circles) across ATP concentrations (mM) expressed as nmol phosphate/mg protein/min. The data is fitted to Michaelis-Menten curves and is presented as mean ± SD (n = 3 biological replicates and n = 6 technical replicates). \**P* = 0.0026 at 0.1 mM ATP; \**P* = 0.0008 at 0.25 mM ATP; \**P* = 0.0006 at 0.5 mM ATP; \**P* = 0.0006 at 1.0 mM ATP; \**P* = 0.0003 at 2.5 mM ATP; \**P* = 0.0009 at 5.0 mM ATP; multiple t-tests. (b) Fold change of ATPase activity at 0.5 mM ATP relative to protein amounts of P-CFTR in LMNG detergent pre-treated with PC, brain PS or a mix of lipids PE/brain PS/egg PC/cholesterol (PE/PS/PC/chol) at 5:2:1:1 weight ratio for 1 h relative to P-CFTR in LMNG detergent. ATPase activity at 0.5 mM ATP of P-CFTR in amphipol was also compared as a reference. The data is presented as mean ± SD (n = 4 biological replicates, n = 4 technical replicates for P-CFTR in LMNG and P-CFTR in LMNG pre-treated with PE/PS/PC/chol; n = 3 biological replicates, n = 3 technical replicates for P-CFTR in LMNG pre-treated with brain PS, PC and P-CFTR in amphipol). ns, not significant; \*\**P* = 0.0012; \*\*\*\**P* < 0.0001; One-way ANOVA with Dunnett’s multiple comparisons test comparing each condition to P-CFTR in LMNG detergent.

In order to determine the role of lipids in enhancing the ATPase activity of the amphipol preparation, we tested the effect of re-introducing PC, PS or a mix of PE:PS:PC:cholesterol (5:2:1:1 ratio by weight) in the LMNG preparation (Figure 4b). We measured a significant increase in ATPase activity of purified P-CFTR in LMNG detergent micelles upon pre-treatment with PE:PS:PC:cholesterol (5:2:1:1 ratio by weight) albeit only reaching half of the relative ATPase activity of purified P-CFTR in amphipol (Figure 4b). These results suggest that these lipids, including cholesterol, enhance catalytic activity of the functional CFTR molecules in the LMNG preparation and that the number of functional CFTR molecules is greater in the amphipol preparation.

We were then prompted to determine if the amphipol preparation enhanced the functional reconstitution of CFTR channel activity relative to the LMNG preparation. Each preparation was separately reconstituted into pre-formed POPC liposomes. We measured anion electrodiffusion from liposomes normalized for the amount of CFTR inserted into liposomes (Refer to *CFTR quantification* section in Materials and Methods for more details on the method) in order to determine the relative proportion of channel competent CFTR molecules from the two preparations (Eckford, Li et al. 2012, Eckford, Li et al. 2015). Briefly, in this assay, CFTR protein in amphipols or LMNG micelles was reconstituted into POPC proteoliposomes with equal concentrations of potassium iodide inside and potassium glutamate outside of the liposomes to provide a driving force for iodide efflux and to maintain the osmolarity (Eckford, Li et al. 2012, Eckford, Li et al. 2015). Addition of valinomycin, an ionophore that selectively facilitates potassium ion flux out of the proteoliposomes, alleviates the charge build up in the proteoliposomes and allows for iodide efflux through activated CFTR (Eckford, Li et al. 2012, Eckford, Li et al. 2015). A schematic of this assay is shown in Figure 5a.

**Figure 5:**
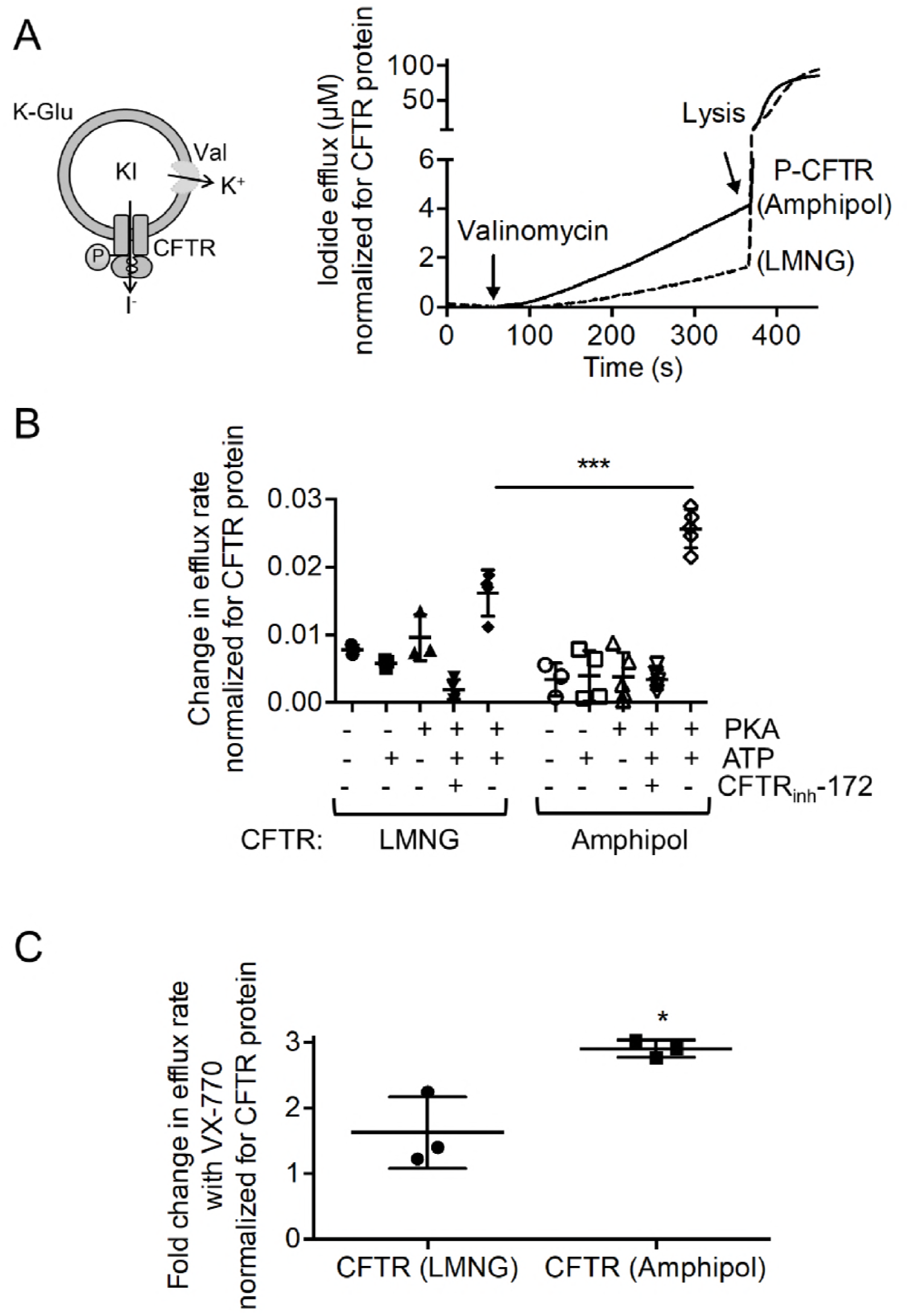
Amphipol protects channel-active conformation of CFTR. (a) Cartoon showing that proteoliposomes are loaded with potassium iodide (KI) on the inside with an equal concentration of potassium glutamate (K-Glu) on the outside. Valinomycin (Val), a potassium ionophore, provides a counterion pathway thereby facilitating iodide (I^-^) electrodiffusion via activated CFTR. The amount of I^-^ effluxed can be detected by an iodide-selective electrode. Representative iodide efflux traces showing iodide effluxed (µM) normalized for CFTR protein (ng) reconstituted in liposomes. Iodide efflux traces of proteoliposomes reconstituted with P-CFTR pre-treated with Mg-ATP in amphipol and of proteoliposomes reconstituted with P-CFTR pre-treated with Mg-ATP in LMNG that were both treated with valinomycin and then lysed with Triton. (b) Change in iodide efflux rate normalized to CFTR amounts (ng) before and after valinomycin treatment of reconstituted P-CFTR in LMNG detergent and in amphipol with Mg-ATP along with negative controls: unphosphorylated CFTR with and without Mg-ATP, P-CFTR without Mg-ATP and P-CFTR with Mg-ATP treated with CFTR_inh_-172. Reconstituted P-CFTR in LMNG detergent and amphipol with Mg-ATP resulted in significant increases in change of slope of iodide efflux compared to negative controls. Reconstituted P-CFTR in amphipol with Mg-ATP resulted in a significantly higher change in slope of iodide efflux compared to reconstituted P-CFTR in LMNG detergent with Mg-ATP. The data is presented as mean ± SD (n = 3 biological replicates, n > 3 technical replicates). \*\*\**P* < 0.0001; One-way ANOVA with Tukey’s multiple comparisons test. (c) Fold change in iodide efflux rate normalized for CFTR protein with 1 µM VX-770 compared to vehicle shows that the potentiation effect of VX-770 was significantly higher in proteoliposomes reconstituted with P-CFTR in amphipol compared to proteoliposomes reconstituted with P-CFTR in LMNG detergent micelles. The data is presented as mean ± SD (n = 3 biological replicates, n = 3 technical replicates). \**P* = 0.0355; paired t-test.

We measured iodide efflux after valinomycin addition to proteoliposomes containing either amphipol or LMNG preparations of CFTR (Figure 5a). As expected for CFTR-mediated flux, electrodiffusion of iodide was minimal for unphosphorylated CFTR or phosphorylated CFTR in the absence of Mg-ATP (Figure 5b) (Eckford, Li et al. 2012). In addition, the CFTR inhibitor, CFTR_inh_-172, reduced flux for the phosphorylated protein in the presence of Mg-ATP, pointing to the specificity of this function (Figure 5b) (Eckford, Li et al. 2012). Together, the proteoliposomal flux assay reports on the phosphorylation and ATP-dependent channel activity of CFTR. Importantly, we observed a significantly higher rate of iodide efflux from proteoliposomes containing P-CFTR purified using amphipol relative to the proteoliposomes containing P-CFTR purified using LMNG (Figure 5b). Given that these proteoliposomal flux studies were controlled for the abundance of total CFTR reconstituted in proteoliposomes and the number of proteoliposomes (estimated by total trapped iodide, Supplementary Figure 5), these findings show that the proportional functional reconstitution is greater in the amphipol preparation. These findings can also be interpreted to suggest that the lipids including cholesterol retained in the CFTR: amphipol complexes exert an activating effect on the CFTR channel.

Ivacaftor (or VX-770) potentiates the regulated channel activity of Wt-CFTR and is approved as a therapeutic intervention for a number of disease-causing mutations (Van Goor, Hadida et al. 2009, Eckford, Li et al. 2012, Jih and Hwang 2013, Van Goor, Yu et al. 2014). To date, its binding site is unknown, although we previously showed using reconstituted CFTR purified using the detergent, *fos*-choline 14, that VX-770 directly modulates channel opening of PKA phosphorylated CFTR (Eckford, Li et al. 2012). Previous studies have suggested that the lipophilic compound, VX-770, interacts with the lipid bilayer and may bind at a CFTR: lipid interface (Jih and Hwang 2013, Baroni, Zegarra-Moran et al. 2014, Hwang, Yeh et al. 2018). Thus, we reasoned that CFTR extracted with its associated lipids using amphipol would exhibit a greater response to VX-770 than CFTR extracted with detergents. Now, we show in paired studies, that the fold increase in iodide electrodiffusion caused by VX-770 (1 μM) from proteoliposomes containing amphipol extracted CFTR was approximately twice that measured for proteoliposomes containing LMNG extracted protein (Figure 5c). We interpret these findings to suggest that the lipids present in the CFTR: amphipol complexes facilitate VX-770 binding and/or the conformational changes required for its potentiation.

## Discussion

We provide direct evidence that lipid interaction with purified CFTR enhances its catalytic function, an activity that is coupled to its ion channel activity. Furthermore, the lipids associated with CFTR participate in VX-770 mediated potentiation, supporting the hypothesis that VX-770 mediates its effects by binding at a protein: lipid interface (Baroni, Zegarra-Moran et al. 2014, Hwang, Yeh et al. 2018).

We also report several methodological advances in the study of CFTR. Our studies suggest that purification of membrane proteins that are relatively intractable, such as CFTR, can be achieved using amphipol A8-35 with yields that are comparable to those achieved by detergents. There are several other examples of membrane proteins that have been directly extracted from membranes by amphipol A8-35 (Popot, Berry et al. 2003, Popot 2018, Marconnet 2019). Further, affinity chromatography can be used to purify membrane proteins either extracted from membranes with amphipol A8-35 (Popot 2018) or after exchanging detergent for amphipol A8-35 (Giusti, Kessler et al. 2015) since the affinity tag is accessible to solution and not embedded in the amphipol belt (Etzkorn, Zoonens et al. 2014). Once purified together with native lipids interacting in a complex with amphipols, these proteins will exhibit enhanced stability and improved suitability for biophysical studies (Marconnet 2019).

This is not the first study to show that CFTR purified in detergents can be functionally reconstituted as a phosphorylated and ATP-regulated anion channel in phospholipid liposomes (Bear, Li et al. 1992, Li, Ramjeesingh et al. 1996, Ramjeesingh, Li et al. 1997, Kogan, Ramjeesingh et al. 2002, Ramjeesingh, Ugwu et al. 2008). In the very first purification papers, we used relatively harsh detergents to extract and purify CFTR prior to reconstitution (Bear, Li et al. 1992, Ramjeesingh, Li et al. 1997, Eckford, Li et al. 2012, Eckford, Li et al. 2015). We estimated that the proportion of functionally reconstituted proteins after detergent purification was low, less than 20% of the total number of CFTR molecules incorporated into liposomes (Bear, Li et al. 1992). Unfortunately, the proportion of purified CFTR that can be functionally reconstituted as an anion channel was not routinely defined in other published purification protocols so a systematic comparison using values in the literature is not possible. However, our proteoliposomal flux studies in the current work show that we can at least double the proportion of channel competent molecules using the amphipol A8-35 relative to the detergent LMNG.

MD simulations carried out on phosphorylated, ATP-bound hCFTR revealed potential phospholipid binding sites. Importantly, this model was built based on a template structure (zCFTR), which shares 55% of sequence identity with hCFTR and shows a nonconductive state. As previously mentioned, preferential binding of POPC was found at the external cleft between TM7 and TM9 and preferential binding of POPS was found at the inward facing gaps between TM4-TM6 and TM10-TM11 (El Hiani and Linsdell 2015, El Hiani, Negoda et al. 2016). It has been speculated that TM8 plays a major role in gating of the conductance path (Zhang and Chen 2016, Liu, Zhang et al. 2017, Zhang, Liu et al. 2017, Hwang, Yeh et al. 2018), hence the localization of a putative POPC interaction site near TM8 (the groove between TM7 and TM9) implicates a role for this lipid in modulation of the gate. Similarly, POPS association with a region in the “pre-open” state of the CFTR channel, that was previously shown to comprise an aqueous portal in cysteine accessibility and electrophysiology studies (El Hiani and Linsdell 2015, El Hiani, Negoda et al. 2016, Hwang, Yeh et al. 2018, Li, Cowley et al. 2018), points to its possible modulation of the channel opening transition. Altogether, the simulations suggest that lipids interact with regions that participate in conformational changes leading to pore opening and suggest that delipidation during purification in detergents may disrupt this mechanism. This interpretation aligns with our functional studies that revealed the importance of lipids in maintaining the conduction competent configuration of CFTR.

We reason that in addition to an important role for the phospholipid interactions modeled in our simulations, the cholesterol identified in the CFTR: amphipol complexes also contributes to the enhanced function mediated by this complex. Cholesterol is present at the plasma membrane and interacts with a population of CFTR in microdomains known as lipid rafts (Cholon, O’Neal et al. 2010, Abu-Arish, Pandzic et al. 2015). The interaction of CFTR and cholesterol regulates the confinement and dynamics of CFTR at the cell surface (Cholon, O’Neal et al. 2010, Abu-Arish, Pandzic et al. 2015). However, not much is known on the effect of the interaction of CFTR and cholesterol on CFTR channel activity. This is the first study to show a direct role for cholesterol in augmenting the intrinsic catalytic and channel activity of CFTR.

We showed previously that the channel activity of detergent purified CFTR was potentiated by ivacaftor (VX-770) following its reconstitution into liposomes, supporting the claim that VX-770 acts via direct binding to CFTR (Eckford, Li et al. 2012). Another group recently studied the interaction of VX-770 with detergent solubilized CFTR using hydrogen/deuterium exchange (Byrnes, Xu et al. 2018). This method showed that there were multiple regions in CFTR that underwent changes in conformation after VX-770 interaction; however, the specific drug binding site has yet to be defined. We and others hypothesize that VX-770 binds at a lipid: CFTR protein interface given its lipophilicity (log*P* score of 5.76) (Baroni, Zegarra-Moran et al. 2014, Chin, Hung et al. 2018) and that VX-770 has been found to insert into the lipid bilayer (Baroni, Zegarra-Moran et al. 2014). In this study, we provide direct evidence that the potentiating effect of VX-770 is facilitated in reconstitutions of CFTR where native lipid interactions are preserved.

In summary, it is clear from these studies of purified and reconstituted CFTR, that the intrinsic catalytic and channel activities of CFTR are modulated by associated lipids. This work points to the need to define the molecular basis for these interactions in order to advance our understanding of channel gating, conduction and the mechanism of action of clinically approved drugs such as ivacaftor (VX-770).

## Materials and Methods

### Homology modeling of human CFTR

A homology model of the NBD-dimerized hCFTR was constructed using MODELLER (Fiser and Sali 2003) with the cryo-EM structure of phosphorylated, ATP-bound *Danio rerio* CFTR channel (zCFTR, PDB: 5W81) as template (Zhang, Liu et al. 2017). The sequence of zCFTR (5W81), denoted as ΔzCFTR below, does not contain segments connecting each TMD-NBD pair (406-436, 1182-1202), the R-domain (642-843), the glycosylated extracellular loop connecting TM7 and TM8 (887-917), and the C-terminal stretch after NBD2 (1459-1485). The full-length sequence of hCFTR and the sequence of PDB 5W81 were aligned with *align2d* program of MODELLER (Zhang, Liu et al. 2017). To avoid modelling long *de novo* segments of zCFTR and prevent short dangling fragments in the final homology model, the protein sequence alignment between hCFTR and ΔzCFTR generated by *align2d* was edited to exclude segments that are not aligned, are too short, or have no corresponding structural information from cryo-EM. The final sequence used for homology modelling consists of five discontinuous segments (hCFTR; 1-403, 438-593, 846-886, 910-1172, and 1204-1457). Fifty homology models of human CFTR were generated using the *model-single* program, and the model with the lowest global DOPE score was chosen as the final model. The resulting structural model of human CFTR was superimposed onto PDB 5W81 with respect to the Cα’s of aligned residues through a translation-rotation operation that minimizes RMSD.

### Preparation of simulation systems

Two Mg-ATP moieties from the zCFTR cryo-EM structure were inserted at the NBD interface of hCFTR by superimposing the hCFTR homology model onto zCFTR using VMD (Humphrey, Dalke et al. 1996). All five proteogenic segments of hCFTR were acetylated at the N-termini and amidated at the C-termini. This hCFTR and Mg-ATP construct was embedded in either POPC or POPS bilayers in hexagonal unit cells. hCFTR was embedded in a POPC bilayer using two InflateGRO methods with slight modification catering to hexagonal geometry (Kandt, Ash et al. 2007, Schmidt and Kandt 2012). This procedure resulted in a bilayer consisting of 306 and 265 lipid molecules for InflateGRO and InflateGRO2 methods, respectively. hCFTR embedded in a POPS bilayer was prepared and solvated using CHARMM-GUI (Jo, Kim et al. 2008), resulting in a bilayer consisting of 287 lipid molecules. To add water and ions to the POPC-embedded systems, tools from GROMACS package were used (Abraham MJ 2015). For POPS-embedded systems, CHARMM-GUI automatically adds water and ions after adding the lipid bilayer. Both POPC-embedded and POPS-embedded systems were solvated in an aqueous solution of 150 mM NaCl in a hexagonal unit cell. No water molecules or ions were placed within the channel pore or within the lipid bilayer in both initial systems. The number of atoms in the simulation cells were 181,000 to 208,000 and unit cell dimensions were: *a* = *b* = 105∼111 Å, *c* = 165∼191 Å, *α* = *β* = 90°, *γ* = 120°.

### MD simulations

The CHARMM36 force field was used for protein, ions, lipids and ATP along with the TIP3P water model (Jorgensen 1983, MacKerell, Bashford et al. 1998, Klauda, Venable et al. 2010, Huang and MacKerell 2013). All systems (POPC-embedded by InflateGRO, POPC-embedded by InflateGRO2, POPS-embedded) were subject to steepest descent energy minimization until the maximum computed forces were below 1000 kJ/mol/nm. All simulations were conducted in the *NpT* ensemble with *T*=300 K, *p*=1 atm, and with a timestep of 2 fs. Constant temperature and pressure were maintained via Nosé–Hoover thermostat (Nosé 1984, Hoover 1985) (temperature coupling of 0.5 ps) and Parrinello-Rahman barostat (Parrinello 1980, Nosé 1983) (time constant of 2.0 ps), respectively. Lennard-Jones interactions were cut off at 1.2 nm and a force-based switching function with a range of 1.0 nm was used. Electrostatic interactions were calculated using particle-mesh Ewald (Darden T 1993, Essmann U 1995) with a real-space cut-off distance of 1.2 nm. Nonbonded interactions were calculated using Verlet neighbor lists (Verlet 1967, Páll 2013). All covalent bonds involving hydrogen atoms were constrained using the LINCS algorithm (Hess 2008). Simulations were performed using GROMACS 2018.1 (Abraham MJ 2015). All systems were *NpT*-equilibrated in three 10 ns-stages before production runs, successively with protein heavy atoms, protein backbone atoms, and protein Cα atoms restrained. All positional restraints used a force constant of 1000 kJ/mol/nm^2^. Random initial velocities were generated for twenty replicas of POPC-embedded system (embedded using either InflateGRO or InflateGRO2 procedures), followed by 1 µs production runs of each replica, for an aggregate total sampling time of 20 µs for POPC-embedded hCFTR. Random initial velocities were generated for ten replicas of the POPS-embedded system (embedded using CHARMM-GUI), followed by 1 µs production runs of each replica, for an aggregate total sampling time of 10 µs for POPS-embedded hCFTR.

To obtain values of bulk lipid densities for normalizing lipid spatial distribution functions of protein-embedded systems, a pure POPC bilayer and a pure POPS bilayer in rectangular cells were constructed and surrounded by explicit water and 150 mM NaCl using CHARMM-GUI membrane builder (cell dimensions: 50 × 50 × 69 Å^3^). The neat bilayer systems contained a total of 84 and 74 lipid molecules for POPS and POPC, respectively. Each system was simulated for 50 ns using GROMACS 2018.1 (Abraham MJ 2015) using identical simulation parameters as those used for protein-embedded bilayer simulations.

### Model validation

The alignment carried out by *align2d* program of the MODELLER gave a different sequence alignment from that reported by Zhang and colleagues in the C-terminal region of NBD1. For example, residues 593-597 (protein sequence: LMASK) of zebrafish CFTR was incorrectly aligned to residues 602-606 (protein sequence: LVTSK) of human CFTR instead of residues 594-598 (protein sequence: LMANK). The differently aligned sequences were deleted from our model because they were regarded as short dangling sequences; as a result, the homology model did not include structure for a small region of NBD1 for which structural information exists in the template structure. To verify whether this truncation alters the conformational and dynamic properties of the transmembrane region of hCFTR, we reproduced all molecular simulations using the original zCFTR model. Molecular models and simulations of zCFTR were conducted using an identical protocol to the methods described above for hCFTR using the structure of zCFTR (PDB 5W81). These simulations were performed for POPS and POPC lipids (the former was embedded using CHARMM-GUI; the latter was embedded using both the InflateGRO and InflateGRO2 protocols), for identical simulation lengths.

To examine structural fluctuations, we computed the RMSD of three subsets of Cα atoms (“All Residues”, “All Without C-terminus”, and “TM Helices”) over simulation time for both the hCFTR homology model and zCFTR, as shown in Supplementary Figure 1. For the “All Residues” subset, all CFTR residues are included except the segments not modeled by cryo-EM (zCFTR) or MODELLER (hCFTR), as explained in the detailed methods of homology modeling. The “All Without C-terminus” subset is the “all” subset minus residues 1438-1485 in zCFTR or 1437-1480 in hCFTR. The C-terminal stretches of both zCFTR and hCFTR are found to be highly dynamic during simulation and account for considerable variation in RMSD amongst the replicas. For the “TM Helices” subset, TM1-TM12 helices and the Lasso motif helices are included. This subset includes all helices that form contacts with lipids. Overall, the RMSDs of zCFTR and hCFTR converged to similar values, especially when only TM helices were considered. The results indicate that the structure of the TM region was not significantly affected by the exclusion of the segment during our simulations. To examine the convergence of average protein-lipid interactions, we computed the average number of lipids within 4.5 Å of protein heavy atoms over simulation time for zCFTR and hCFTR (Supplementary Figure 2). The average number of protein-lipid interactions are stable throughout simulations and zCFTR and hCFTR models are within one standard deviation for the majority of simulation time.

### Analysis of MD simulations

Prior to analysis, all simulation frames were aligned using the Cα atoms of all TM segments of the cryo-EM structure. Analysis was performed on all simulation frames spaced by 5.0 ns after removing the first 100 ns of data from each simulation repeat. For time-averaged analyses, only POPC-embedded by InflateGRO2 and POPS-embedded simulations were utilized based on the agreement we observed between zCFTR and hCFTR models (Supplementary Figure 1). Protein-lipid contact propensities were measured for each amino acid of hCFTR by computing a binary contact (yes/no) whenever any protein-lipid heavy-atom pair was less than 4.5 Å using MDTraj (McGibbon, Beauchamp et al. 2015). These data were calculated for each simulation repeat and averaged over all repeats.

Spatial distribution functions for all lipids were calculated as a number density on a grid with 1 Å spacing using MDAnalysis (Michaud-Agrawal, Denning et al. 2011). POPC and POPS spatial distribution functions (g_PC(r)_ and g_PS(r)_) were normalized with respect bulk lipid number density in a neat bilayer system. PC and PS spatial distribution functions were computed using all simulation frames and the average number density of the bilayer was computed (by truncating all six edges of the box along X, Y, and Z by 10 Å, and taking a mean of the entire lipid region, both lipid headgroups and tails). The number average lipid density was computed to be 0.0956 ± 0.0152 and 0.1015 ± 0.0138 Å^-3^ for PC and PS, respectively. POPC and POPS spatial distribution functions were normalized by dividing by the bulk values PC and PS in the neat bilayer systems. Volume maps were rendered at an isovalue cutoff of 1.6 for POPC and POPS; showing regions with > 60% increased density relative to the bulk bilayer concentration. Difference maps were computed by computing the difference between spatial distribution functions (g_PC(r)_ - g_PS(r)_, and g_PS(r)_ - g_PC(r)_). Volumes of the difference map were rendered at an isovalue cutoff of 0.8; showing regions with an absolute difference > 80% between PC and PS, and vice versa. Manipulation of spatial distribution functions was performed using the volutil functionality of VMD (Humphrey, Dalke et al. 1996).

### Generation of FLAG-CFTR-His construct

N-terminal DYKDDDDK (FLAG) tag was introduced on the CFTR-His construct (with a C-terminal 10x His-tag) on the plasmid DNA containing WT-CFTR cDNA (in pcDNA3.1) as the template using the KAPA HiFi HotStart Ready Mix (KAPA Biosystems, Wilmington, MA), primers (Forward: 5’-GAG ATG GAT TAT AAA GAT GAT GAT G-3’, Reverse: 5’-CAT CAT CAT CTT TAT AAT CCA TCT C-3’) and polymerase chain reaction. The construct was validated by DNA sequencing (TCAG, Toronto, Ontario, Canada) and amplified by transformation and plasmid maxi-prep (Qiagen, Hilden, Germany). High-quality (260/280 nm ratio of 1.8 and higher) plasmid DNA was concentrated to a final concentration of 1 μg/μl.

### Expression of FLAG-CFTR-His construct in HEK293F cells and generation of crude membranes

The FLAG-CFTR-His construct was transiently transfected in HEK293F (Thermo Fisher Scientific) suspension cell line capable of complex N-glycosylation. DNA at 50 μg was mixed in a 1:1 ratio with transfection reagent FectoPRO^®^ (Polyplus, Berkeley, CA) and added to each 200 ml suspension culture at 0.8 x 10^6^ cells per ml, as previously described (Ereno-Orbea, Sicard et al. 2017, Ereno-Orbea, Sicard et al. 2018). Cells were grown at 37°C shaking at 180 rpm with 8% CO_2_ for 24 h. The next day, cells were treated with 10 mM sodium butyrate (Sigma-Aldrich, St. Louis, MO) at 37°C shaking for 24 h. HEK293F cells transfected with FLAG-CFTR-His were collected and spun down at 6000 rpm at 4°C for 30 min. Fresh or frozen cell pellet generated from a 600 ml cell suspension was resuspended in 40 ml of phosphate-buffered saline (PBS, Wisent Inc., Saint-Jean Baptiste, Quebec, Canada) containing one tablet of cOmplete, EDTA free protease inhibitor (Roche, Mannheim, Germany). The cells were lysed using Emulsiflex C3 high pressure homogenizer (Avestin, Ottawa, Ontario, Canada) with at least 5 passages during the lysing process. Cell debris were spun down at 1000g at 4°C for 15 min. The supernatant from this spin was further spun down at 100,000g at 4°C for 2 h. The pelleted crude membrane was resuspended in 20 ml of PBS containing 0.1 mM DDM (BioShop, Burlington, ON, Canada) and protease inhibitor (Roche). Syringes with 21G and 27G needles (BD, Franklin Lakes, NJ) were used to disaggregate the pellet. The DDM treated membranes were incubated on ice for 10 min. Any heavy particulate matter that came down were discarded before spinning the membrane suspension at 100,000g at 4°C for 2 h. The extracted crude membrane pellet was again resuspended with the aid of the syringe and previously described needles in 20 ml of 25 mM HEPES, 100 mM NaCl buffer with protease inhibitor (Roche) and then aliquoted. Total protein concentration of the crude membranes was determined by Bradford assay (Bradford 1976) and presence of CFTR was confirmed by immunoblotting.

### Purification of FLAG-CFTR-His in amphipol or detergent

Crude membranes were solubilized to a final concentration of 2 mg/ml with 1% LMNG (Anatrace, Maumee, OH) and 0.2% cholesterol hemisuccinate (Anatrace) in buffer A (20 mM Tris-HCl pH 7.5, 2 mM MgCl_2_, 200 mM NaCl, 20% glycerol, and 2 mM DTT, with protease inhibitor, Roche) as previously described (Zhang and Chen 2016, Liu, Zhang et al. 2017, Zhang, Liu et al. 2017) or with 3% (w/v) amphipol A8-35 (Anatrace) in buffer B (25 mM HEPES pH 8, 100 mM NaCl with protease inhibitor, Roche) at 4°C for 2 h with shaking. Insoluble fractions were removed by ultracentrifugation at 45,000 rpm at 4°C for 1 h. Anti-FLAG^®^ M2 magnetic beads (Sigma-Aldrich) were washed with corresponding buffers (buffer A with 0.025% LMNG or buffer B with 0.025% amphipol) using a magnetic rack (New England Biolabs, Ipswich, MA) for a total of 3 times. Soluble fractions were incubated with anti-FLAG^®^ M2 magnetic beads at 4°C overnight with shaking. Unbound proteins were removed the next day with a magnetic rack (New England Biolabs) and anti-FLAG^®^ M2 magnetic beads (Sigma-Aldrich) were washed 5 times with corresponding buffers. If phosphorylation is required, the anti-FLAG^®^ M2 magnetic beads (Sigma-Aldrich) were treated with a phosphorylation cocktail (10,000 U of PKA (New England Biolabs) and 5 mM Mg-ATP (Sigma-Aldrich) in buffer A with 0.025% LMNG or buffer B with 0.025% amphipol) at room temperature for 1 h with shaking. The phosphorylation cocktail was removed by magnetic rack and the anti-FLAG^®^ M2 magnetic beads were washed again for 5 times with the corresponding buffers. FLAG-CFTR-His protein was eluted with 200 μg/ml FLAG^®^ peptide (DYKDDDDK, Sigma-Aldrich) in buffer A with 0.025% LMNG or buffer B with 0.025% amphipol using the magnetic rack after 45 min incubation with shaking at 4°C. Purified protein at 500 μl was loaded in Superose 6 Increase 10/300 column (GE Healthcare, Chicago, IL) to further characterize the protein. Purity of purified CFTR was determined by densitometry of the CFTR bands compared to other bands present on the 4-12% silver stained gels by Image Studio™ Lite software.

### CFTR quantification

Crude membranes at 2.6 mg were solubilized in amphipol and CFTR was purified in amphipol as previously described to generate 400 μl of eluted standard in the presence of FLAG^®^ peptide. An aliquot of standard (100 μl) was subjected to amino acid analysis with the Waters Pico-Tag System (Waters ACQUITY UPLC, Milford, MA). Refer to Supplementary Figure 3 and Supplementary Table 1 for complete results of amino acid analysis of the purified CFTR standard in amphipol. Alanine (1015.112 pmoles was present in the 100 μl standard) was chosen to determine the concentration of CFTR standard as amino acid hydrolysis with various solvents resulted in 100% recovery of alanine (Adebiyi, Jin et al. 2005) and there are no alanine residues in the FLAG^®^ peptide that was also present in the standard. The calculation of the concentration of the CFTR standard is as follows:

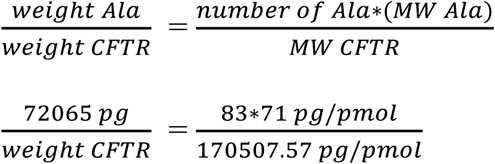

Weight of CFTR = 208512269 pg or 2085.12269 ng

Volume used for amino acid analysis = 100 µl

Concentration of CFTR standard = 2085.12269 ng / 100 µl = 20.85 ng/µl

The remaining standard was aliquoted and stored at −80°C. When required, various nanogram amounts of the standard were run with purified protein samples of unknown amounts on SDS-PAGE and subjected to immunoblotting. Immunoblot bands were analyzed with Image Studio™ Lite software and densitometry of unknowns were interpolated on the standard curve on GraphPad Prism. Refer to Supplementary Figure 4 for example of standard curve for concentration determination.

### Phospholipid assay

Purified CFTR in amphipol, purified CFTR in LMNG and the appropriate controls were subjected to a 96 well phospholipid assay kit according to the manufacturer’s protocol (Cat. no. MAK122, Sigma-Aldrich). Fluorescence was read at excitation of 530 nm and emission of 585 nm on the SpectraMax i3X plate reader (Molecular Devices, San Jose, CA).

### Lipid TLC

Lipid containing samples were extracted as previously described (Dorr, Koorengevel et al. 2014). The lipid containing organic phase was first dried with sodium sulfate for 1 h, filtered and concentrated under a stream of argon to a volume of 50 μl. Silica gel plates (Analtech Inc., Newark, DE) were first activated at 160°C for 1 h before sample application. Samples (10 μl) and polar lipid standard (2 μl, Matreya LLC, State College, PA) were spotted and developed using chloroform: methanol: water (65:25:4) as the solvent system. Visualization of the lipids was conducted as previously described (Dorr, Koorengevel et al. 2014).

### ATPase assay - ATP dose response

Purified protein (approximately 50-200 ng in 25 μl) was incubated with 50 μl of various concentrations of Na-ATP (Sigma-Aldrich) solubilized in 10 mM HEPES, 100 mM NaCl, 5 mM MgCl_2_, 1 mM DDM, pH 7.4 buffer at 37°C for 2 h. A colorimetric assay was applied as previously described (Chifflet, Torriglia et al. 1988) to detect phosphate at absorbance 800 nm on the SpectraMax i3X plate reader (Molecular Devices). Values were subtracted from controls (amphipol or LMNG buffer with various concentrations of ATP), converted to nmol phosphate by interpolation of phosphate standard curve on GraphPad Prism and then to nmol phosphate/mg protein/min after CFTR quantification as previously described. Values were fit on Michaelis-Menten curves with GraphPad Prism to determine the K_m_ and V_max_ values of ATPase activity.

### ATPase assay - Re-introducing lipids to purified CFTR in LMNG detergent micelles

Egg PC, porcine brain PS, or a mix of PE/brain PS/egg PC/cholesterol in a 5:2:1:1 (w/v) ratio (all lipids in chloroform from Avanti^®^ Polar Lipids, Inc., Alabaster, AL) were dried in a glass tube under argon gas. Dried lipids were resuspended in 25 mM HEPES, 100 mM NaCl, pH 8.0 buffer with sonication to a stock concentration of 10 mg/ml. Resuspended lipid (100 μg in 10 μl) was added to purified protein (approximately 50-200 ng in 25 μl) on a 96 well plate along with controls: 1) 10 μl of 25 mM HEPES, 100 mM NaCl, pH 8.0 buffer added to 25 μl of purified protein; 2) 10 μl of resuspended lipid added to 25 μl of buffer A with 0.025% LMNG; 3) 10 μl of 25 mM HEPES, 100 mM NaCl, pH 8.0 buffer added to 25 μl buffer A with 0.025% LMNG. Protein with lipids and controls were incubated at 1 h at room temperature. Na-ATP (40 μl of 0.9375 mM stock concentration to a final concentration of 0.5 mM Na-ATP) solubilized in 10 mM HEPES, 100 mM NaCl, 5 mM MgCl_2_, 1 mM DDM, pH 7.4 buffer was added to the mix and the reaction was incubated at 37°C for 2 h. Colorimetric assay was applied as previously described above.

### Iodide efflux

CFTR protein in amphipol was incubated with 5 mM DDM during the elution step with FLAG^®^ peptide prior to reconstitution whereas protein in LMNG remained as is. Purified protein (approximately 0.4 μg) was reconstituted in 5 mg of POPC (Avanti^®^ Polar Lipids, Inc.) and subjected to iodide efflux as previously described (Eckford, Li et al. 2012, Eckford, Li et al. 2015).

## Acknowledgements

We would like to thank Dr. Manuela Zoonens for comments on the manuscript. We would also like to thank Dr. Jacqueline McCormack for help in editing the manuscript. We would like to thank Dr. Steven Molinski for help in generating the FLAG-CFTR-His construct as well as Yu-Sheng Wu for help and support throughout the project. We would like to thank Rey Interior of the SPARC BioCentre at the Hospital for Sick Children for conducting the HPLC for amino acid analysis.

## Author contributions

Study was designed by S.C. and C.E.B. based on discussions with the other authors. S.C. generated and analyzed the SEC, CFTR quantification, phospholipid, ATPase and iodide efflux data with the help from M.R., J.E.-O., C.E.B. and J-P.J. M.R. assisted in optimization of the novel purification method, ATPase assay, lipid TLC, and amino acid quantification of CFTR standard. M.H. assisted in optimization of the novel purification method. J.E.-O. conducted transfections of CFTR in HEK-293F cells in suspension and conducted SEC of purified protein. H.C. conducted transfections of CFTR in HEK-293F cells. C.I. and Z.W.Z. generated homology model of hCFTR, and conducted and analyzed MD simulations. R.P. provided suggestions on presentation and interpretation of MD simulation data. J-P.J assisted with optimization of novel purification method and suggestions for improvement of assays. Manuscript was written by S.C. and C.E.B. with input from all the authors.

## Supplementary Figures and Tables Legends

**Supplementary Figure 1:**
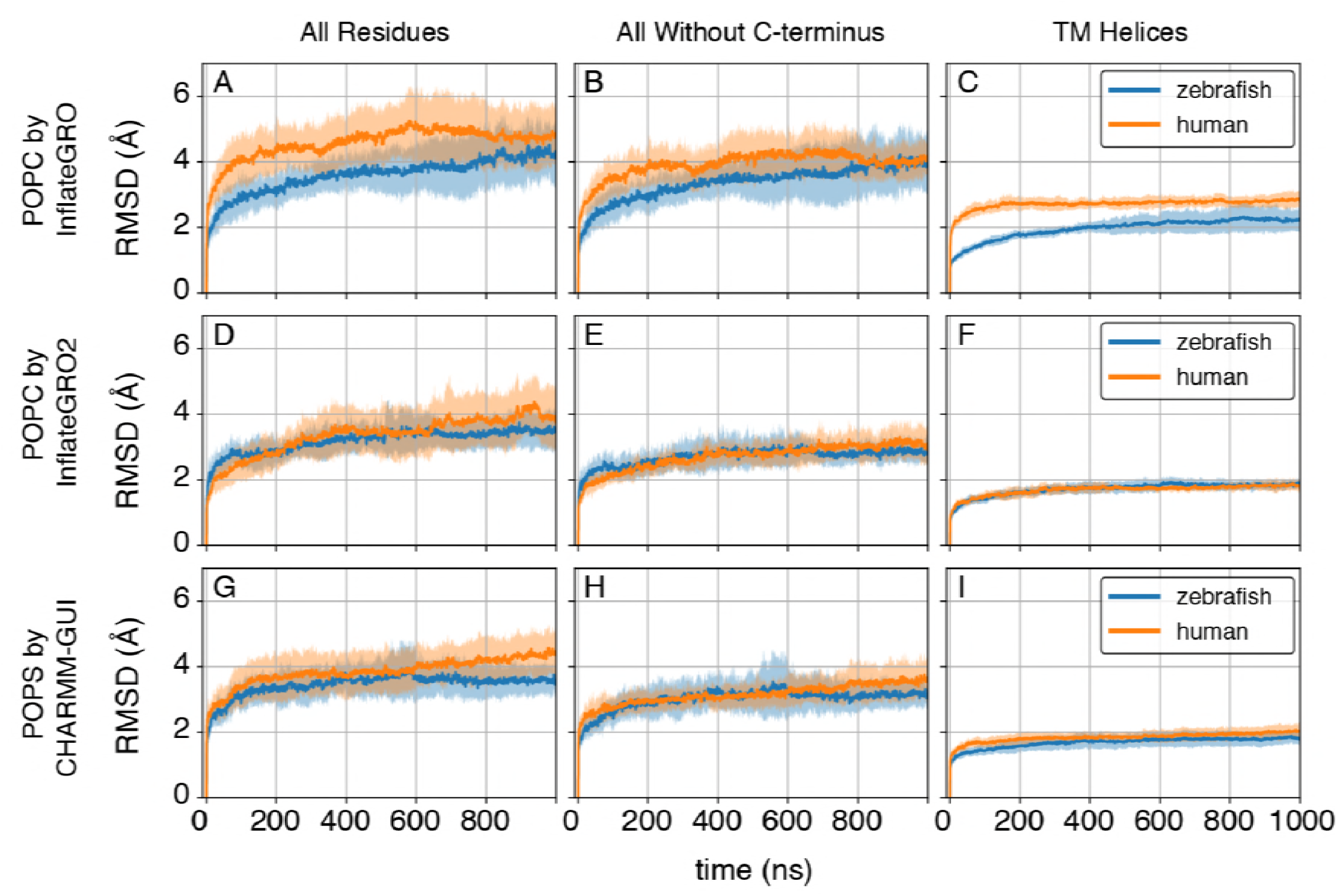
Cross-validation of hCFTR and zCFTR models. RMSD of Cα atoms versus time, averaged over simulation replicas. Regions within one standard deviation from the mean RMSD are shaded. Each row identifies a bilayer type and embedding method; (a-b-c) POPC-embedded using InflateGRO, (d-e-f) POPC-embedded using InflateGRO2, (g-h-i) POPS-embedded using CHARMM-GUI. The three columns show RMSD values with respect to the initial structures for different subsets of protein Cα atoms for hCFTR (orange) and zCFTR (blue) models. Both zCFTR and hCFTR RMSD values are within one standard deviation of the mean for systems POPC-embedded using InflateGRO2 and POPS-embedded using CHARMM-GUI.

**Supplementary Figure 2:**
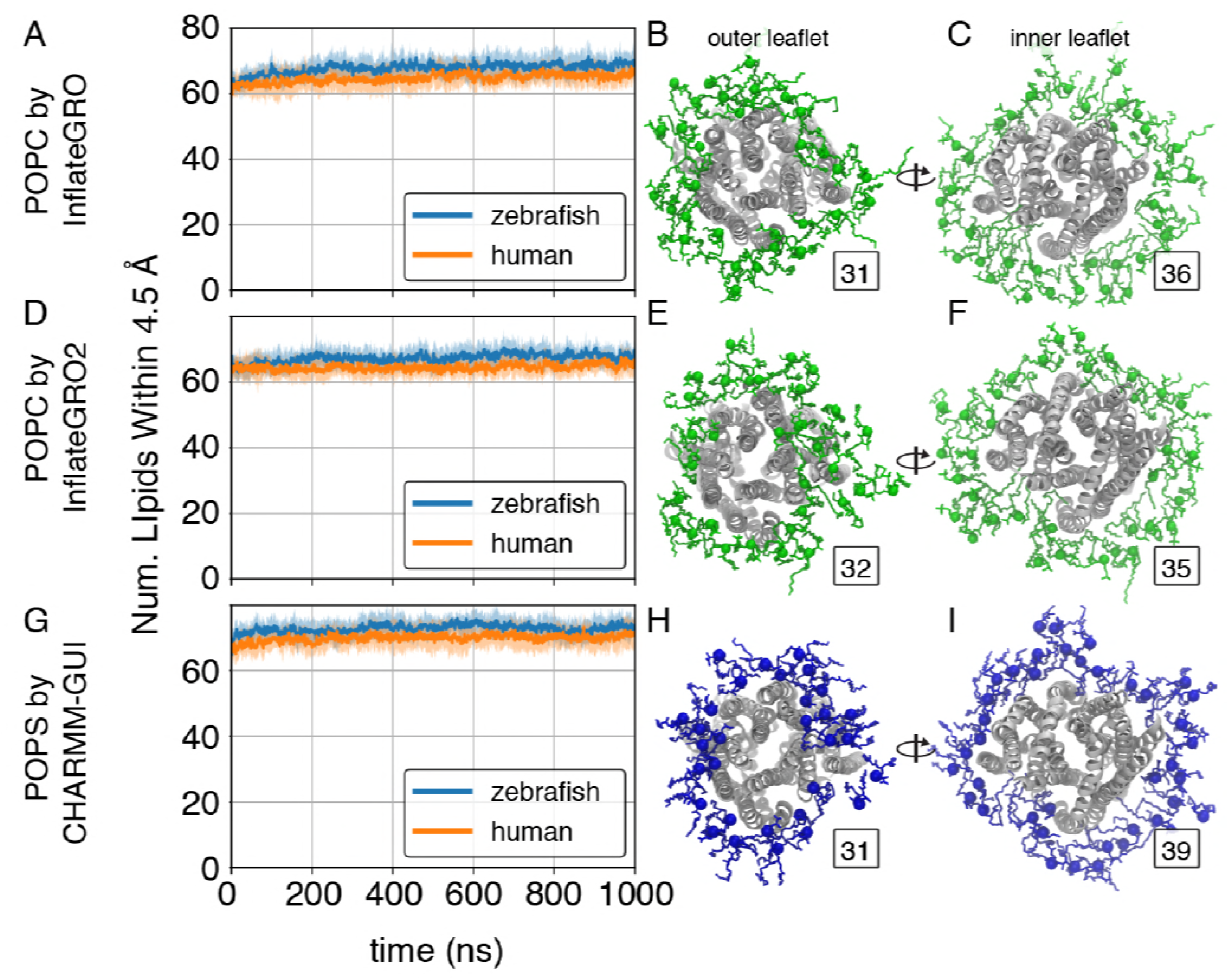
Protein-lipid interactions in hCFTR and zCFTR models. Average number of lipids interacting with protein heavy atoms versus time. All lipid molecules with a heavy-atom pair distance less 4.5 Å are counted. Regions within one standard deviation from the mean are shaded. Each row corresponds to a bilayer type and embedding method; (a) POPC-embedded using InflateGRO, (d) POPC-embedded using InflateGRO2, (g) POPS-embedded using CHARMM-GUI. In these panels, average lipid counts from simulations conducted with the hCFTR (orange) and zCFTR (blue) models are shown. For each bilayer type and embedding method, molecular renderings of individual snapshots depicting all lipid molecules within 4.5 Å of protein heavy-atoms are shown from the extracellular and intracellular orientation (b-c, e-f, h-i). POPC (green) and POPS (blue) are shown in licorice representation with phosphorous headgroup atoms depicted as spheres. The total number of lipids in the outer and inner leaflet within 4.5 Å are shown as an inset number.

**Supplementary Figure 3:**
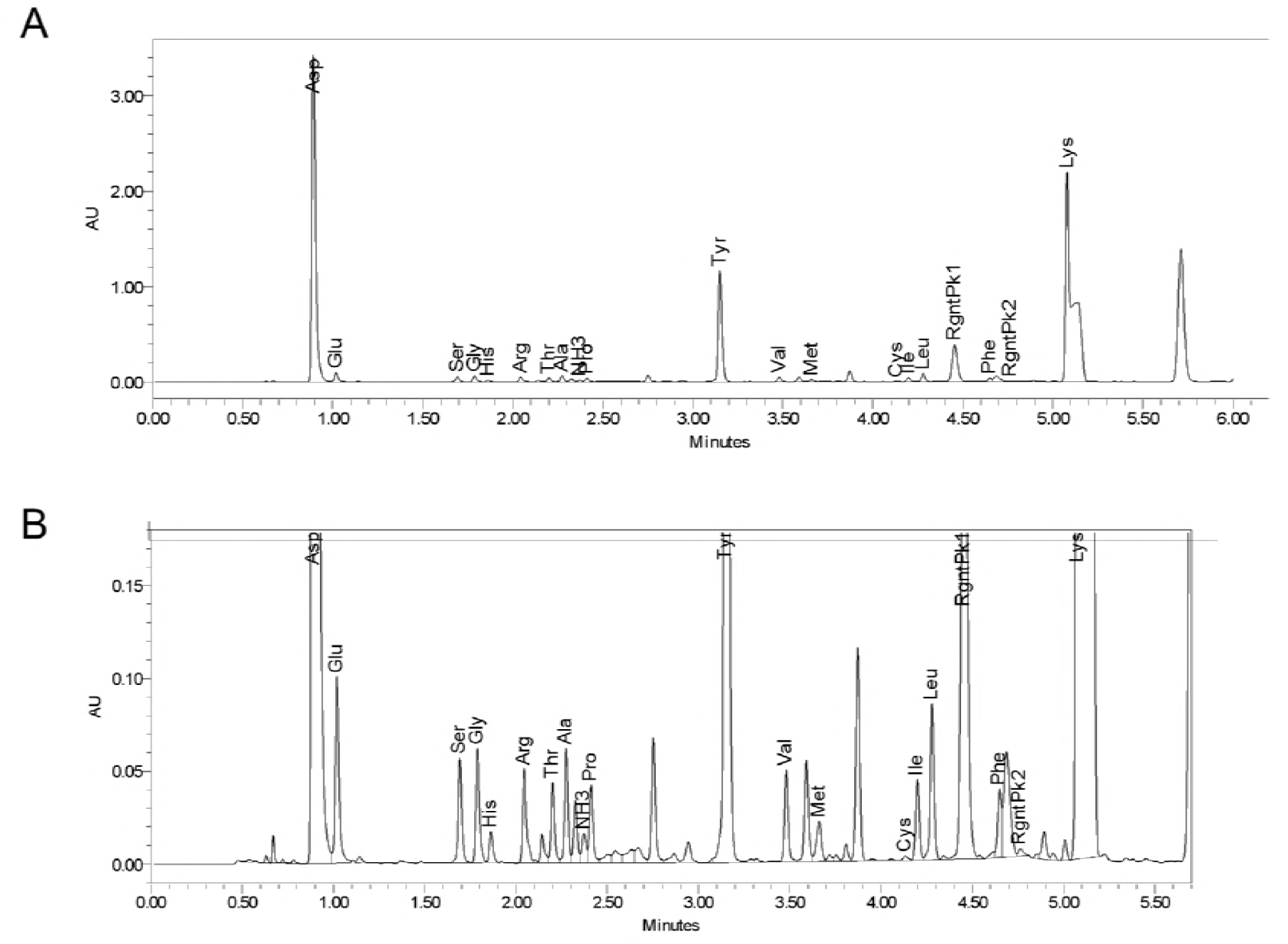
Results of amino acid analysis of CFTR standard. (a) Chromatogram of all the amino acids present in CFTR standard. (b) Scaled chromatogram to magnify the smaller peaks.

**Supplementary Figure 4:**
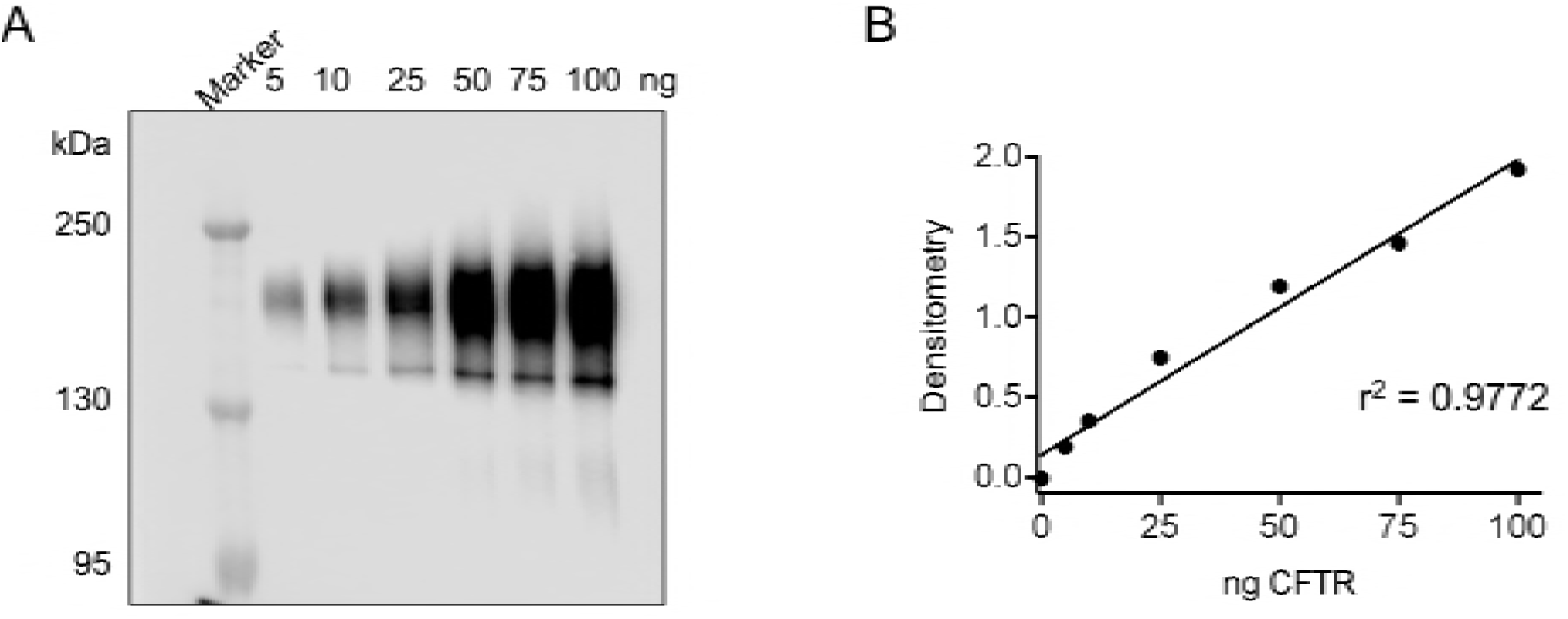
Representative immunoblot and standard curve applied for CFTR quantification. (a) Immunoblot of CFTR standard at increasing amounts (nanograms). (b) Standard curve of (a) that is used for interpolation of unknown purified protein.

**Supplementary Figure 5:**
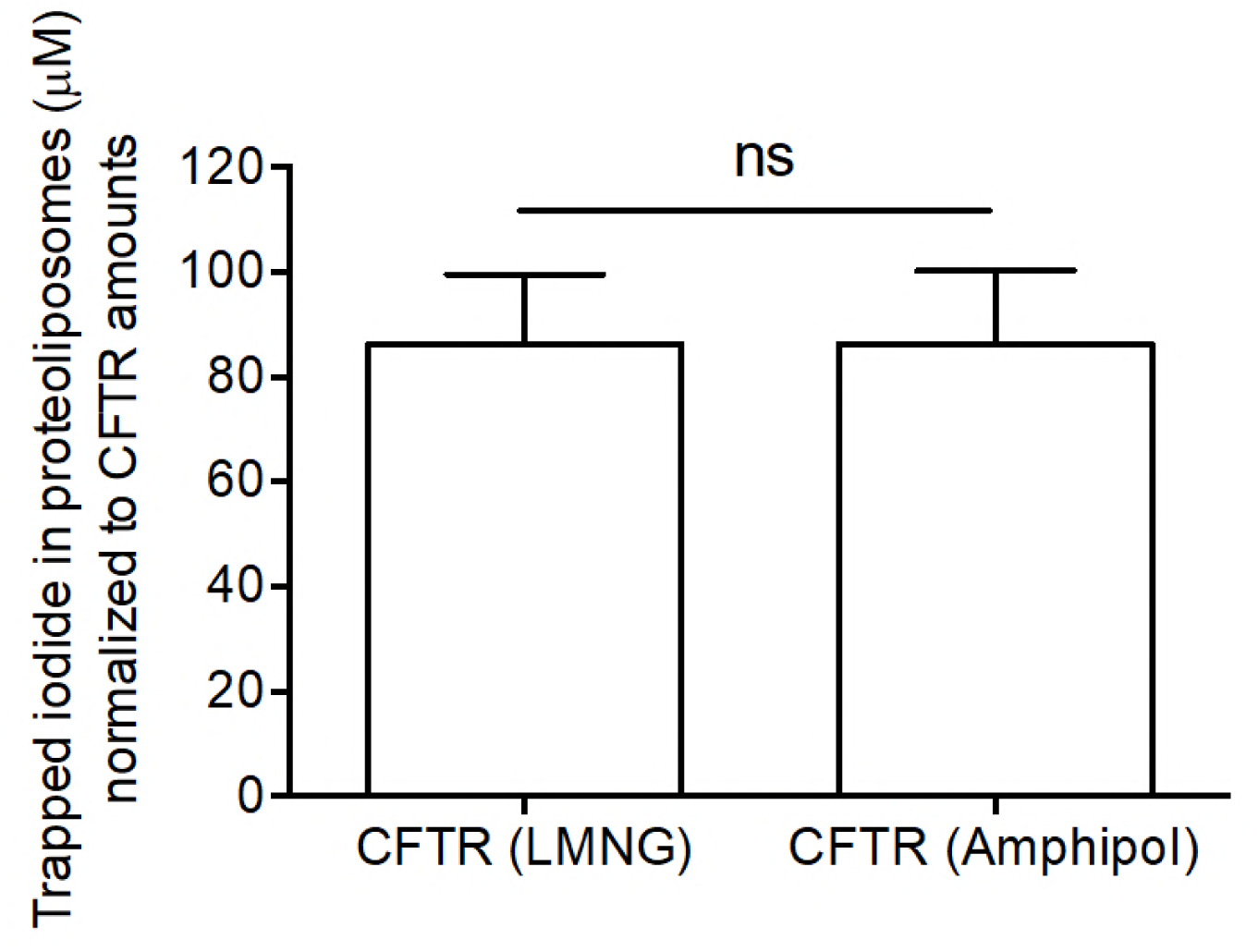
Trapped iodide is similar for the two reconstitutions, reconstitution of LMNG purified CFTR and amphipol purified CFTR. Bars correspond to iodide measurements before valinomycin addition and after detergent lysis normalized to CFTR amounts. The data is presented as mean ± SD (n > 3 biological replicates and n > 3 technical replicates). *P* = 0.9974; paired t-test.

**Supplementary Table 1:**
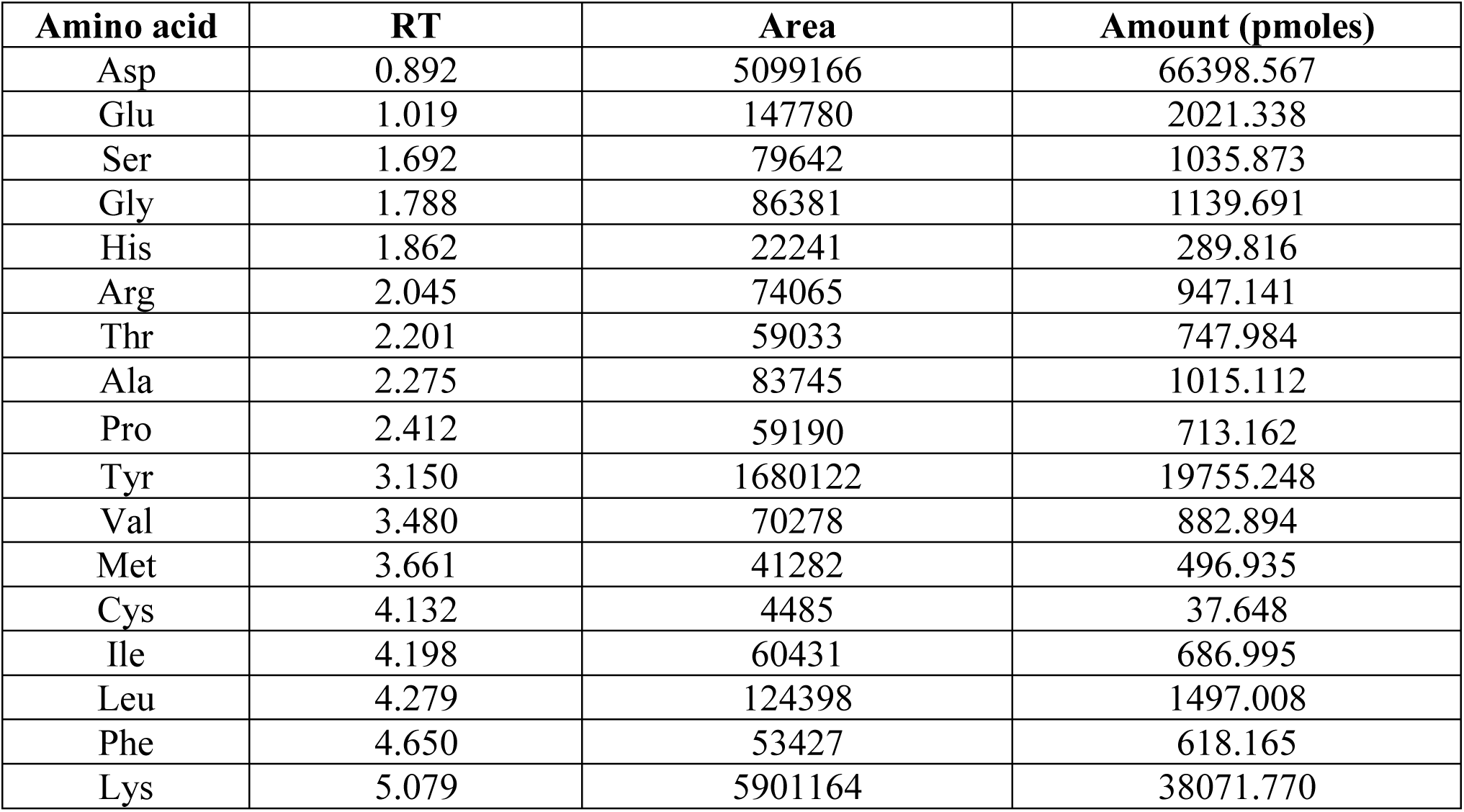
Summary table of amino acid analysis results of CFTR standard. Table shows the amino acid (3-letter code), retention time (RT) in minutes of when the amino acid shows up, the area under the peak and the corresponding amount of the amino acid in picomoles (pmoles) in a 100 µl purified CFTR standard.

**Supplementary Table 2:** Variables for determining concentration of CFTR standard.

| Variable | Value |
| --- | --- |
| MW Ala | 89.0935 – 18 (for peptide bond)<br>= 71 g/mol or pg/pmole |
| weight Ala | 1015.12 pmoles (from chromatogram)<br>1015.12 pmoles x 71.0935 pg/pmole<br>= 72065 pg |
| number of Ala in CFTR | 83 |

